# Lipid droplets 3D full measurement by holographic in-flow tomography

**DOI:** 10.1101/2021.12.09.471789

**Authors:** Daniele Pirone, Daniele Sirico, Lisa Miccio, Vittorio Bianco, Martina Mugnano, Danila del Giudice, Gianandrea Pasquinelli, Sabrina Valente, Silvia Lemma, Luisa Iommarini, Ivana Kurelac, Pasquale Memmolo, Pietro Ferraro

## Abstract

The most recent discoveries in the biochemical field are highlighting the increasingly important role of lipid droplets (LDs) in several regulatory mechanisms in living cells. LDs are dynamic organelles and therefore their complete characterization in terms of number, size, spatial positioning and relative distribution in the cell volume can shed light on the roles played by LDs. Until now, fluorescence microscopy and transmission electron microscopy are assessed as the gold standard methods for identifying LDs due to their high sensitivity and specificity. However, such methods generally only provide 2D assays and partial measurements. Furthermore, both can be destructive and with low productivity, thus limiting analysis of large cell numbers in a sample. Here we demonstrate for the first time the capability of 3D visualization and the full LD characterization in high-throughput with a tomographic phase-contrast flow-cytometer, by using ovarian cancer cells and monocyte cell lines as models. A strategy for retrieving significant parameters on spatial correlations and LD 3D positioning inside each cell volume is reported. The information gathered by this new method could allow more in depth understanding and lead to new discoveries on how LDs are correlated to cellular functions.

## Introduction

Lipid droplets (LDs) are ubiquitous intracellular organelles specialized in triacylglycerols and steryl esters storage, found in some prokaryotes and in most eukaryotic cells, where they reside primarily in the cytoplasm (1). Initially described exclusively as storage organelles, LDs are now recognized as dynamic entities that play several other pivotal roles in intracellular homeostasis. For example, LDs provide a defence mechanism against numerous stress conditions (e.g., lipotoxicity, endoplasmic reticulum (ER) stress, oxidative stress, mitochondrial damage during autophagy), and control certain proteins’ expression by supporting their maturation, storage and turn-over (2, 3). In addition to ER from which they derive, LDs dynamically interact with most intracellular organelles, including mitochondria, peroxisome, lysosomes, Golgi apparatus, and nuclei (2), which contribute to the final three-dimensional (3D) spatial organization of LDs inside the cellular volume. Although the mechanisms linking specific LD structural characteristics to a certain function are still not completely understood, a vast amount of evidences shows that variation in LD number, size, ultrastructure, motility, lipid/protein content and interactions with other organelles significantly influences many cellular processes (2, 4–6). Consequently, their dysregulation may have implications in diseases, and evaluation of LD-related parameters may be exploited as a biomarker. Indeed, LDs have been described to have a role in various pathologies, including diabetes (7), atherosclerosis (8), fatty liver disease (9), pathogen infections (10), neurodegenerative diseases (11) and cancer (12–15). Moreover, they are recognized as structural markers of inflammation, since a remarkable increase in LD number and size rapidly occurs in immune cells in response to inflammatory stimuli (5, 10, 16–18). Most recent evidence shows that monocytes from COVID-19 affected patients display an increased LD accumulation with respect to healthy blood donors (19), suggesting a possible involvement of these organelles in the SARS-CoV-2 pathogenesis (20). Noteworthy, the presence of LDs inside the cell volume has been recently exploited for a completely new purpose, i.e LDs behave as endogenous biological micro-lenses enhancing fluorescence imaging of cellular sub-structures (21). However, the limitations of currently available techniques for LD characterization still prevent from completely understanding their functions and exploiting their potential for clinical purposes. In particular, development of new non-destructive techniques is required, which provide fast LD detection, quantification, and characterization, ensuring powerful statistical data, with the aim of discovering novel insights on this prominent issue. Among the various techniques used for LD investigation, transmission electron microscopy (TEM) and fluorescent microscopy (FM) are probably the most exploited for this purpose (22, 23). The LD ultrastructure can be easily determined by TEM due to their homogeneous spherical shape and their recognizable electron density (24). However, only small areas of a sample can be analysed by TEM, thus strongly limiting the ensemble study of LDs inside the cell, and the method requires skilled and highly trained operators. FM is a somewhat more user-friendly technique, with a growing number of fluorescent lipophilic dyes used for LDs detection, most popular being Nile Red, Bodipy^®^ 493/503, LipidTOX and Oli Red O (25, 26). These reagents come with the advantage of being easy to use, thus allowing tracing LDs dynamics, even by the live cell imaging (26). These dyes may also be used in flow-cytometry, allowing higher throughput, but providing no information on LD spatial distribution. It is important to note that fluorescent dyes are subjected to photobleaching, may interfere with cell function, especially during long exposure times, and induce phototoxicity. These limitations have prompted the development of label-free methods for live imaging.

Among label-free approaches, digital holography (DH) in microscope configuration is a valuable non-destructive tool for LD analysis, as their refractive index (RI) substantially differs from the surrounding cell inner structure, and the DH method relies on the RI difference as a contrast agent. The first demonstration in visualizing and measuring LDs in live cells by DH was reported in the last decade (27). Recently, the tomographic phase-contrast microscopy (TPM) was applied for LD 3D imaging within mammalian cells (28, 29) and microalgae (30), for 4D tracking of the LDs dynamics in live hepatocytes (31), and for recording time-lapses of living foam cells (32). Nevertheless, the up-to-date available label-free techniques allow to investigate LDs only in static, adherent cells, strongly limiting both throughput and reliability of the information regarding the LD spatial organization, the latter being significantly affected by the cell culture mode. Furthermore, imaging methods developed for operating on adhesion samples exclude the possibility to investigate populations that naturally exert their functions in circulation, such as cells in bodily fluids. Although TPM based on DH apparatus are really powerful (33), they cannot furnish in simple way high-throughput analysis that can instead be achieved only by imaging techniques capable to operate in flow-cytometry modality. Instead, an imaging modality for phenotyping the cells in flow-through is highly demanded in order to investigate the sample in an environment that well mimics physiological conditions and can guarantee statistically significant assays by investigating a large number of cells in a single experiment. Recently, a novel approach based on DH-TPM operating in-flow modality has been developed, thus showing great potentialities for achieving quantitative 3D phase-contrast tomograms of red blood cells and diatoms (34), tumor cells (35, 36), and very lately also for 3D visualization and display of internalized graphene nanoparticles in flowing cells (37). Although DH has already been extensively investigated for the two-dimensional (2D) high-throughput analysis of LDs (38), to the best of our knowledge the label-free TPM has never been implemented in-flow cytometry mode to the same scope. The use of in-flow TPM can offer new insights from the imaging and full-characterization of the LDs in 3D for each single cell.

Here we demonstrate for the first time that LDs can be visualized and quantitatively measured in 3D in live cell suspensions while they are flowing along a simple and commercially available microfluidic channel. The presented approach, based on label-free TPM, allows to retrieve, besides all the 2D microscopy-based morphological features, 3D and quantitative information on LDs locations. Numerical image processing is implemented for retrieving the phenotype of each detected LD, their spatial correlations and positioning into the 3D cell volume. To prove the robustness and reliability of this new tomographic microscopy mode, we experimentally investigated the human ovarian cancer cell line A2780 and the monocyte cell line THP-1, and compared our analyses with the state-of-the-art techniques used for LD imaging, i.e. the TEM and FM. The reported results show that a new avenue for achieving high-throughput investigation of the presence and the distribution of intracellular LDs in flowing samples can be achieved. This makes viable the development of novel diagnostic or therapeutical tools in biomedicine capable to furnish exact 3D location of the LDs inside the cells, their volume, shape, RI-based statistics and dry mass at the single-cell level, which cannot be accessed by conventional TEM and FM imaging.

## Results

### Experiments with in-flow TPM

In this section we show the 3D visualization of LDs inside A2780 and THP-1 cells (the number of cells were 54 and 34, respectively) obtained through the 3D label-free in-flow TPM system, and we report an assay of measured LDs parameters. To this aim, cells were introduced inside a microfluidic channel observed through a DH microscopy apparatus. After having recorded the digital hologram video of flowing cells, a numerical processing pipeline has been implemented in order to retrieve the corresponding quantitative phase maps (QPMs). In a QPM, the 3D spatial distribution of the sample’s RIs is coupled to the sample’s physical thickness in the form of a 2D image. To decouple these two information, multiple QPMs of the same sample are needed, which can be obtained by illuminating it along different viewing angles. For this reason, the flux of the cells along the microfluidic channel has been engineered in order to guarantee that the cells rotate while they are passing through the field of view (FOV) of the DH imaging system. Consequently, hundreds of QPMs for each flowing cell, each corresponding to a different viewing angle were obtained. Moreover, the large FOV allows the simultaneous recording of multiple cells, thus contributing to the high-throughput property. After having estimated the viewing/rolling angles, the 3D RI spatial distribution at the single-cell level can be reconstructed. A more detailed description of the in-flow TPM system and the numerical processing for the tomographic reconstruction is reported in the Materials and Methods section.

In this way, the label-free 3D RI tomograms of 54 A2780 live cells have been computed. In Fig. 1A, we show the central slice of the 3D reconstructed tomogram of one typical A2780 cell. The LDs presence is clearly visible. Moreover, it can be noted from the colour scale of the plot that LDs reach RI values much higher than the surrounding medium. In fact, the corresponding RI histogram in Fig. 1B goes up to 1.500, that is a very high value for these types of cells, thus suggesting that a threshold-based method is enough for numerical segmenting the LDs and thus to extract the quantitative measurement of each LD. In Figs. 1C-E we report three isolevels representations of the same 3D tomogram after having segmented LDs with three different thresholds numerical value (i.e., 1.400, 1.420, and 1.440, respectively). All three results are plausible, since separate particles have been isolated in the same cell location. Obviously, the greater the RI threshold, the smaller the volume of particles identified. Hence, a criterion to set the LDs-threshold is requested. The RI values of LDs change based on the type of cell, the temperature, and the wavelength (39). Segmenting intracellular organelles is always problematic in a label-free technique since an exogenous calibrated marker is missed. In the 2D case, a deep learning approach has been employed to identify LDs inside the QPMs (40). Instead, in the 3D case, to segment LDs in microalgal cells (32) and in foam cells (30), the RI threshold has been selected according to the FM image of the same cell obtained from the channel mounted on the static TPM system. However, this is not possible in TPM flow-cytometry. Instead, to fix an average RI threshold, here we exploited independent 2D FM measurements about the number and the diameter of LDs (see the Conventional 2D Imaging section), and the high number of cells reconstructed through the in-flow TPM technique. The TPM configuration selected for these experiments has a lower spatial resolution than the 2D FM images in order to provide a very large FOV (see the Discussion section). Therefore, as shown in Figs. 1C-E, in this experiment LDs cannot be resolved when they are too close each other. For this reason, the number of LDs measured through the in-flow TPM is expected to be smaller than the FM technique. However, the overall volume of LDs must be unchanged. Within the 2D FM images, the average LDs volume per cell can be computed indirectly by multiplying the average number of LDs and their average diameter, thus obtaining 6.10 μm3 in the A2780 case. Instead, as reported in Fig. 1F, the LDs average volume per cell can be measured directly from the 54 tomograms by varying the RI threshold, thus obtaining the expected decreasing curve. The LDs-threshold can be finally selected in such a way to guarantee the same average volume, which is RI ≥ 1.423 in the A2780 case. By using the computed LDs-threshold, all the 54 tomograms of the A2780 cells have been segmented. In Figs. 1G-I, we display three of these segmented tomograms by painting separated LDs or LDs clusters with different colors, and in Figs. 1J-L we report the corresponding RI distributions in 3D, which are very similar to each other.

**Fig. 1.**
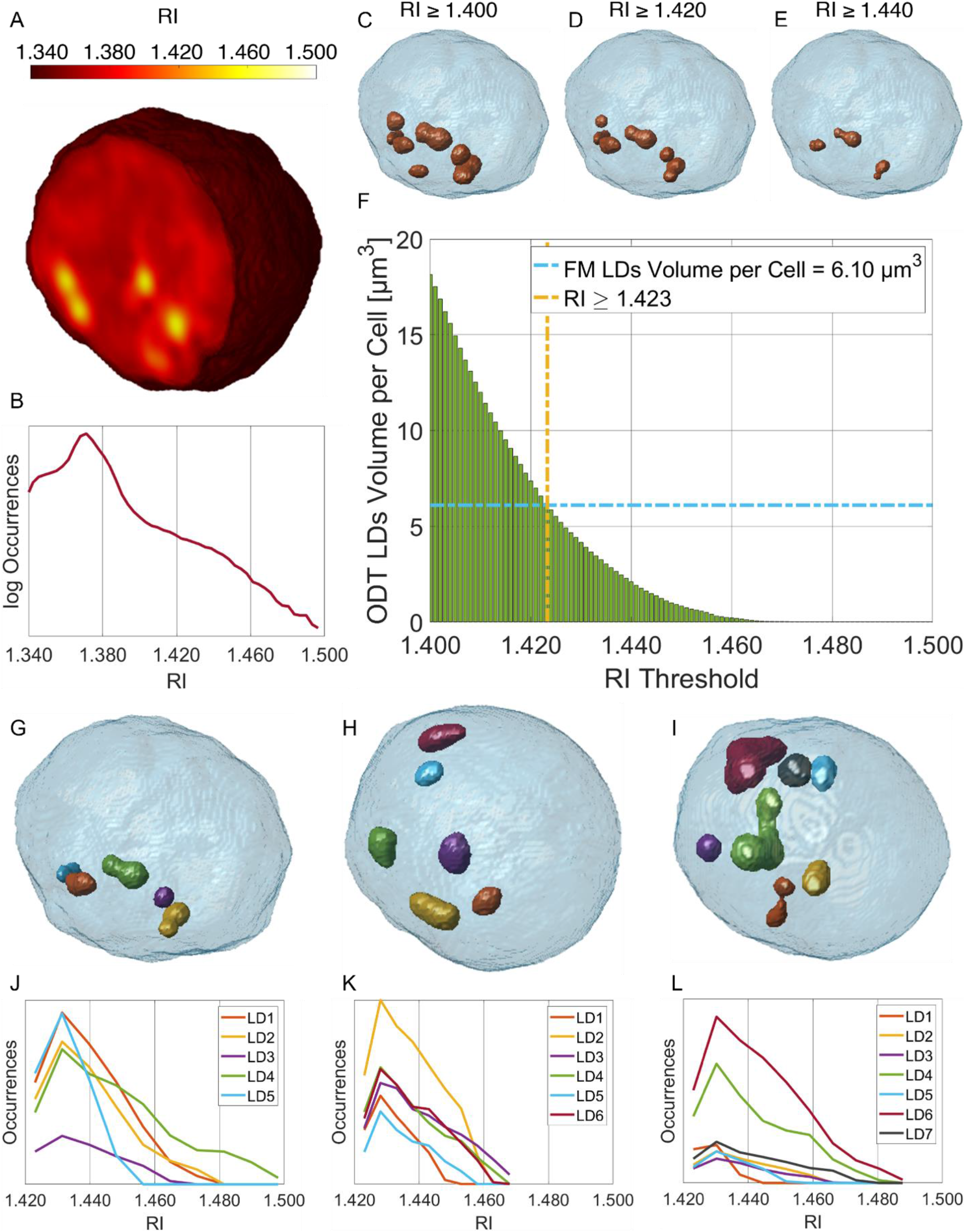
Segmentation of the LDs within the 3D RI tomograms of A2780 live cells. (A) Central slice of the 3D RI tomogram of an A2780 live cell, in which LDs take the highest RI values. (B) Histogram in logarithmic scale of the 3D RI distribution of the cell in (A). (C-E) Isolevels representation of the tomogram in (A), in which LDs (orange) have been segmented by using the RI thresholds reported above. (F) Average volume per cell of the LDs segmented in 54 TPM tomograms of A2780 live cells by using different RI thresholds. The selected LDs-threshold (yellow line) allows computing the same average volume measured in 2D FM images (blue line). (G-I) Isolevels representation of separated LDs or LDs clusters segmented in 3 A2780 tomograms by using the LDs-threshold selected in (F), and (J-L) corresponding RI histograms. (A-E,G,J) are the same cell.

In recent years, TPM has emerged because it allows label-free quantitative measurements at the single-cell level of features about both the 3D morphology and the RI statistics, which are related to the cell biophysical properties (e.g., dry mass) (41). The implementation of the tomographic phase-microscopy system in flow-cytometry environment further allows the replication of the same measurement on a large number of cells, thus reaching a statistical significance which can be exploited for characterizing a certain phenomenon. Therefore, in Fig. 2 we report the histograms of several properties about the hundreds of reconstructed LDs. In particular, in Fig. 2A E we show respectively the mean value, the standard deviation, the entropy, the kurtosis, and the skewness of the 3D RI distribution about each LD. The equivalent radius displayed in Fig. 2F is the radius of a sphere having the same volume of the analyzed LD. The dry mass reported in Fig. 2G is the mass of the biological sample without its water content, calculated as in (28). The sphericity shown in Fig. 2H is instead computed as the ratio between the surface area of a sphere with same volume of the LD and its actual surface area, thus providing a quantification of the particle’s shape (sphericity is 1 if the particle is a perfect sphere, otherwise it is smaller than 1 the more the particle has a non-spherical shape).

**Fig. 2.**
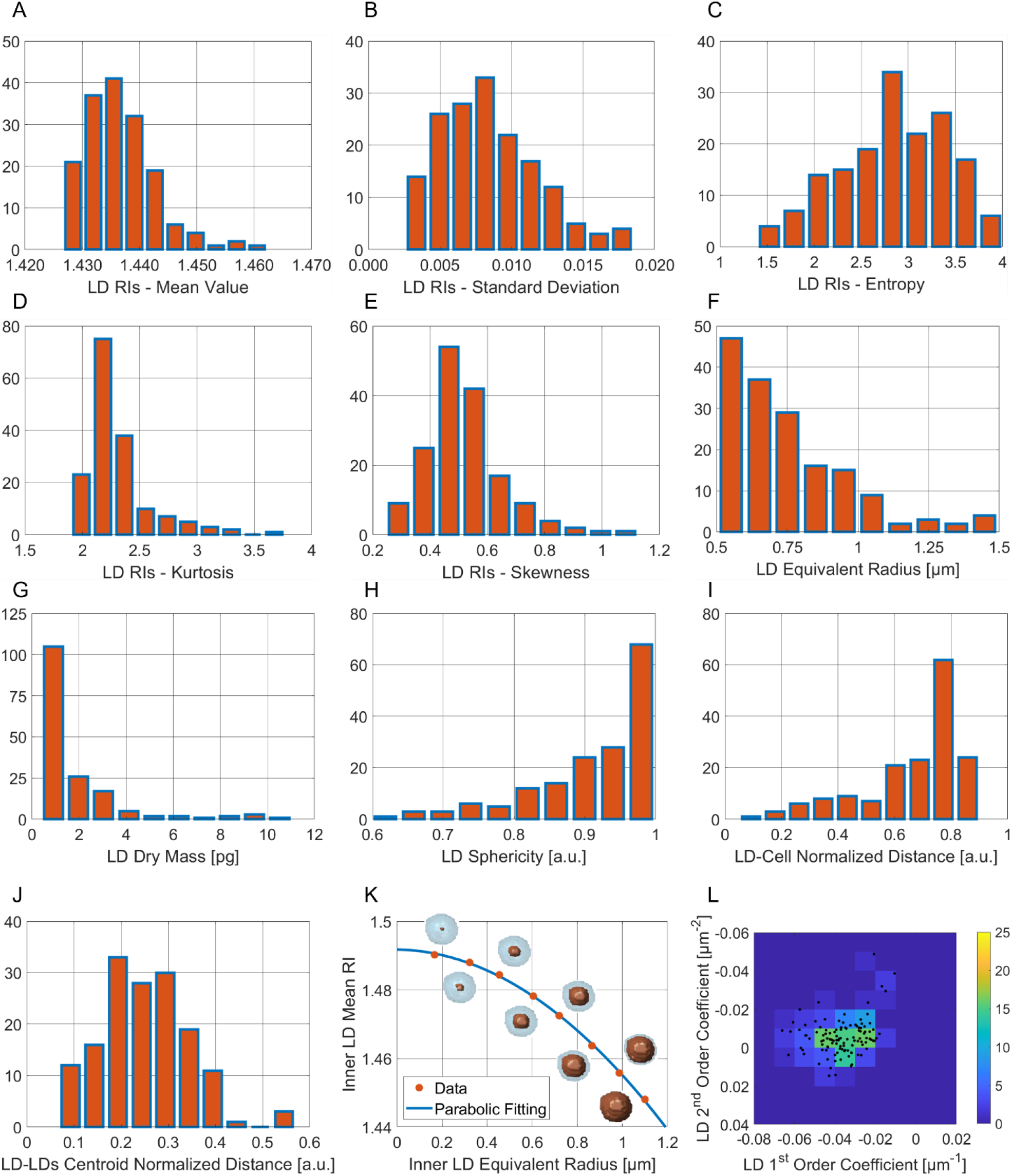
LDs features extracted from 54 A2780 3D RI tomograms. (A-E) Histograms of respectively the mean value, standard deviation, entropy, kurtosis, and skewness of the 3D RI distributions of each LD. (F-H) Histograms of respectively the equivalent radius, the dry mass, and the sphericity of each LD. (I) Histogram of the distance between each LD centroid and the corresponding cell centroid, normalized to the cell equivalent radius. (J) Histogram of the distance between each LD centroid and the centroid of all the LDs inside the same corresponding cell, normalized to the cell equivalent diameter. (K) Mean RIs (orange dots) of concentric inner zones (orange regions) selected inside the same LD, with overlapped in blue the parabolic fitting. (I) Bivariate histogram of the first and second order coefficients of the parabolic fitting in (K) measured in all the LDs (black dots).

Furthermore, the proposed in-flow TPM allows reconstructing the 3D tomograms of suspended cells rather than adhered cells, therefore the 3D spatial arrangement of LDs inside the cell can be accessed and investigated. In particular, in order to set parameters about this assay, we computed the distance between the centroid of each LD and the centroid of the cell that contains it, normalized to the cell equivalent radius. It is worth to remark that the corresponding histogram, reported in Fig. 2I, shows a bimodal distribution. The central region of a cancer cell is usually occupied by the nucleus (42), and LDs are usually expected to be found inside the cytoplasm (43). For this reason, most of LDs are about the 80 % of the cell equivalent radius away from the cell centroid. However, there is a minor amount of LDs nearer to the cell centroid, which can be explained considering that LDs are sometimes found also within the nucleus (44). Moreover, the 3D tomograms in Figs. 1G-I have confirmed the property of LDs of concentrating in the same region of the cytoplasm (45), which has been also observed in the 2D FM images. To quantify this property, for each cell we computed the centroid of all the LDs, and we calculated its distance from each LD, normalized to the cell equivalent diameter. The corresponding histogram is displayed in Fig. 2J, which provides a characterization of the spread of the LDs positions around their own ensemble centroid.

From a structural point of view, LDs are formed by an inner core which mainly stores triacylglycerols and steryl esters and are surrounded by a phospholipid monolayer studded with LD-specific proteins (46, 47). Therefore, the RI is expected to change passing from the outer zone to the inner zone of the LD. For this reason, we considered concentric volumes inside the same LD. In Fig. 2K we show a sequence of the same LD as the size of the internal structure decreases (orange regions) along with the corresponding mean RIs (orange dots). The mean RI increases passing from the overall volume, made of both the membrane and the inner core, to the sole inner core. Moreover, the computed data are perfectly fitted by a parabolic curve. Therefore, we performed the parabolic fitting for all the LDs, and we report their first order and second order coefficients in Fig. 2L (black dots), overlapped to the corresponding bivariate histogram. This analysis confirms the higher density of the inner LD core with respect to its surrounding region. In general, the plots shown in Fig. 2K,L can be used as a tool to inspect the inner LD RI distribution.

By using the same pipeline, we also analyzed about two hundred of THP-1 cells (48). As shown in Fig. 3A, by comparing the average LDs volume per cell with the 2D FM measurement, we selected the LDs-threshold RI ≥ 1.411. This threshold allowed identifying LDs in 34 THP-1 live cells. In Figs. 3B,C, the QPMs of a monocyte without LDs and with LDs are respectively shown, while the central slices of the corresponding 3D RI tomograms are reported in Figs. 3D,E, respectively. In both 2D and 3D cases, the LDs are clearly recognizable as distinguishable spots with the highest phase or RI values, respectively. For this reason, in the histogram of the 3D RI distribution in Fig. 3F, the two cells can be easily identified. Indeed, as expected, only the monocyte with LDs, which isolevels representation is displayed in Fig. 3G, shows an inflated distribution of RIs, due to a large number of occurrences for higher RI values. The 3D RI tomograms of the 34 THP-1 monocytes have been exploited to measure the same features about LDs described in the case of A2780 cells, as shown in Fig. 4. The average values and the standard deviations of these parameters are resumed in Table S1 for both the A2780 and THP-1 cells. On average, LDs in A2780 cells have a greater RI mean value, standard deviation, entropy, kurtosis, and skewness than the THP-1 monocytes. Moreover, as they are bigger in size too, they also have a greater dry mass. Again, the histogram of the LD-cell normalized distance shows a bimodal distribution, see Fig. 4J. In general, the 3D disposition inside the cell is about the same in both cases.

**Fig. 3.**
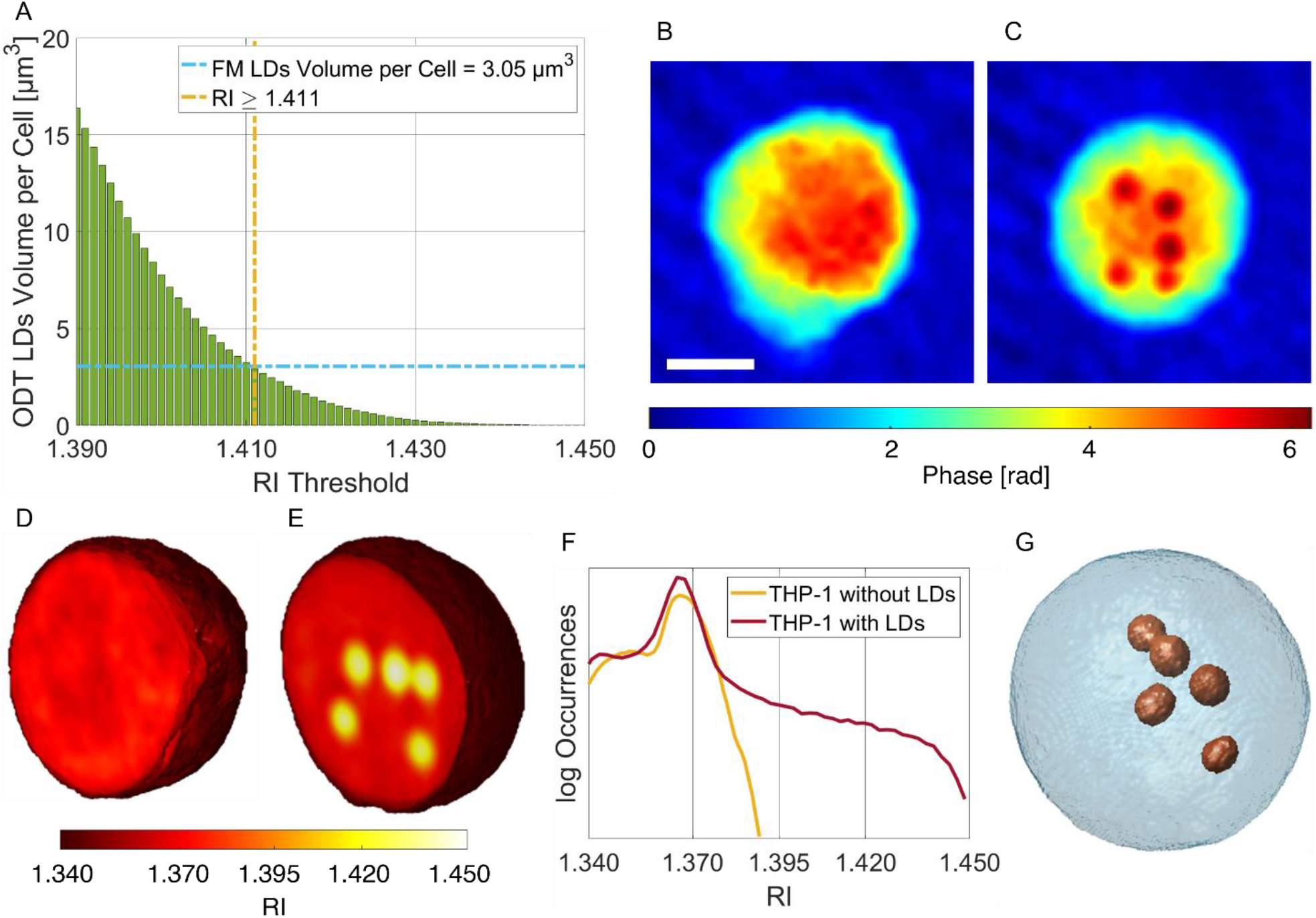
Segmentation of the LDs within the 3D RI tomograms of THP-1 live cells. (A) Average volume per cell of the LDs segmented in 34 TPM tomograms of THP-1 live cells by using different RI thresholds. The selected LDs-threshold (yellow line) allows computing the same average volume measured in 2D FM images (blue line). (B,C) QPMs of two THP-1 cells, one without LDs (B) and the other one with LDs (C) (dark red spots). Scale bar is 5 μm. (D,E) Central slices of the 3D RI tomograms of the cells in (B,C), respectively, in which LDs take the highest RI values. (F) Histogram in logarithmic scale of the 3D RI distribution of the cells in (D) (yellow) and (E) (red). (G) Isolevels representation of the tomogram in (E), in which LDs (orange) have been segmented by using the LDs-threshold selected in (A).

**Fig. 4.**
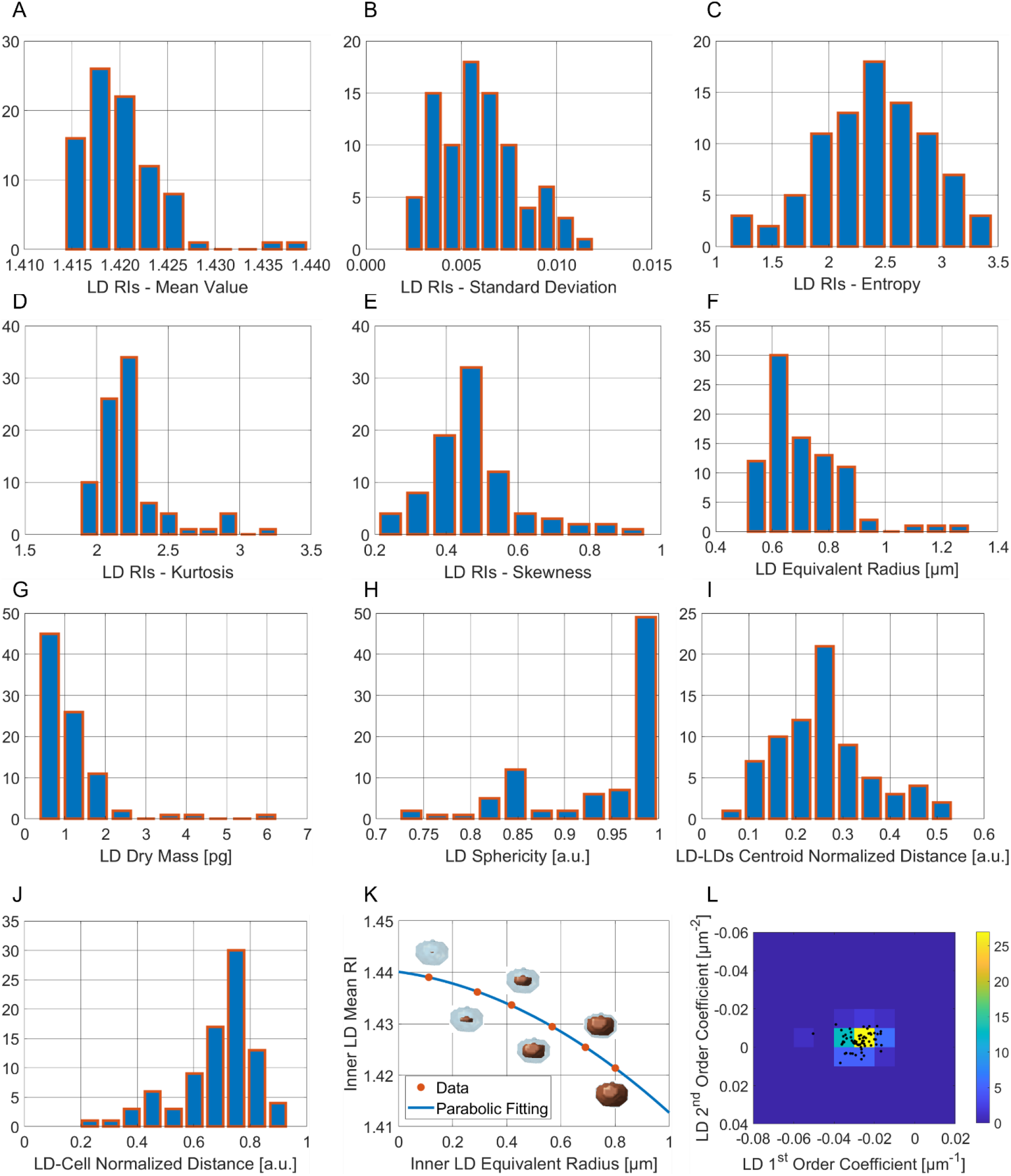
LDs features extracted from 34 THP-1 3D RI tomograms. (A-E) Histograms of respectively the mean value, standard deviation, entropy, kurtosis, and skewness of the 3D RI distributions of each LD. (F-H) Histograms of respectively the equivalent radius, the dry mass, and the sphericity of each LD. (I) Histogram of the distance between each LD centroid and the corresponding cell centroid, normalized to the cell equivalent radius. (J) Histogram of the distance between each LD centroid and the centroid of all the LDs inside the same corresponding cell, normalized to the cell equivalent diameter. (K) Mean RIs (orange dots) of concentric inner zones (orange regions) selected inside the same LD, with overlapped in blue the parabolic fitting. (L) Bivariate histogram of the first and second order coefficients of the parabolic fitting in (K) measured in all the LDs (black dots).

### Conventional 2D imaging

With the aim to demonstrate the advantages of in-flow TPM approach, we next compare our results with the currently available gold-standard techniques for LD analysis, namely TEM and FM upon Nile Red staining. Thanks to the high spatial resolution and the contrast of TEM images, LDs were visible with a high level of details in A2780 and THP 1 cells (Figs. 5A,G), with average LD dimension being 963 nm (SD+/− 242) and 736 nm (SD+/− 150), respectively. The reliable LD counting is not possible when using TEM, due the small FOV and the limited sample section (80 nm) with respect to the cell size (tens of microns). These intrinsic drawbacks of TEM prevent the precise determination of the number of cells harbouring LDs, allowing only approximate estimation. On the other hand, the FM analysis permitted LD quantification, revealing a higher number of LDs in ovarian cancer model (p<0.005, Figs. 5B-F and Figs. 5H-L). In particular, LDs were counted in 11 A2780 and THP-1 live cells, obtaining on average 26.55 (SD+/− 4.18) and 10.55 (SD+/− 1.96) organelles per cell, respectively (Figs. 5E,K). In line with TPM data, LDs were detected in all A2780 cells while they were missing in some THP-1 cells. No significant difference in LD size was observed between the two models. The diameter of 30 LDs per cell type was measured, revealing the average values of 760 nm (SD+/− 39) in A2780 and 820 nm (SD+/− 29) in THP-1 model (Figs. 5F,L). Interestingly, FM images showed that LDs within the A2780 cytoplasm were not uniformly distributed, but mainly assembled near the nucleus, confirming the same phenomenon observed in the TPM reconstructions.

**Fig. 5.**
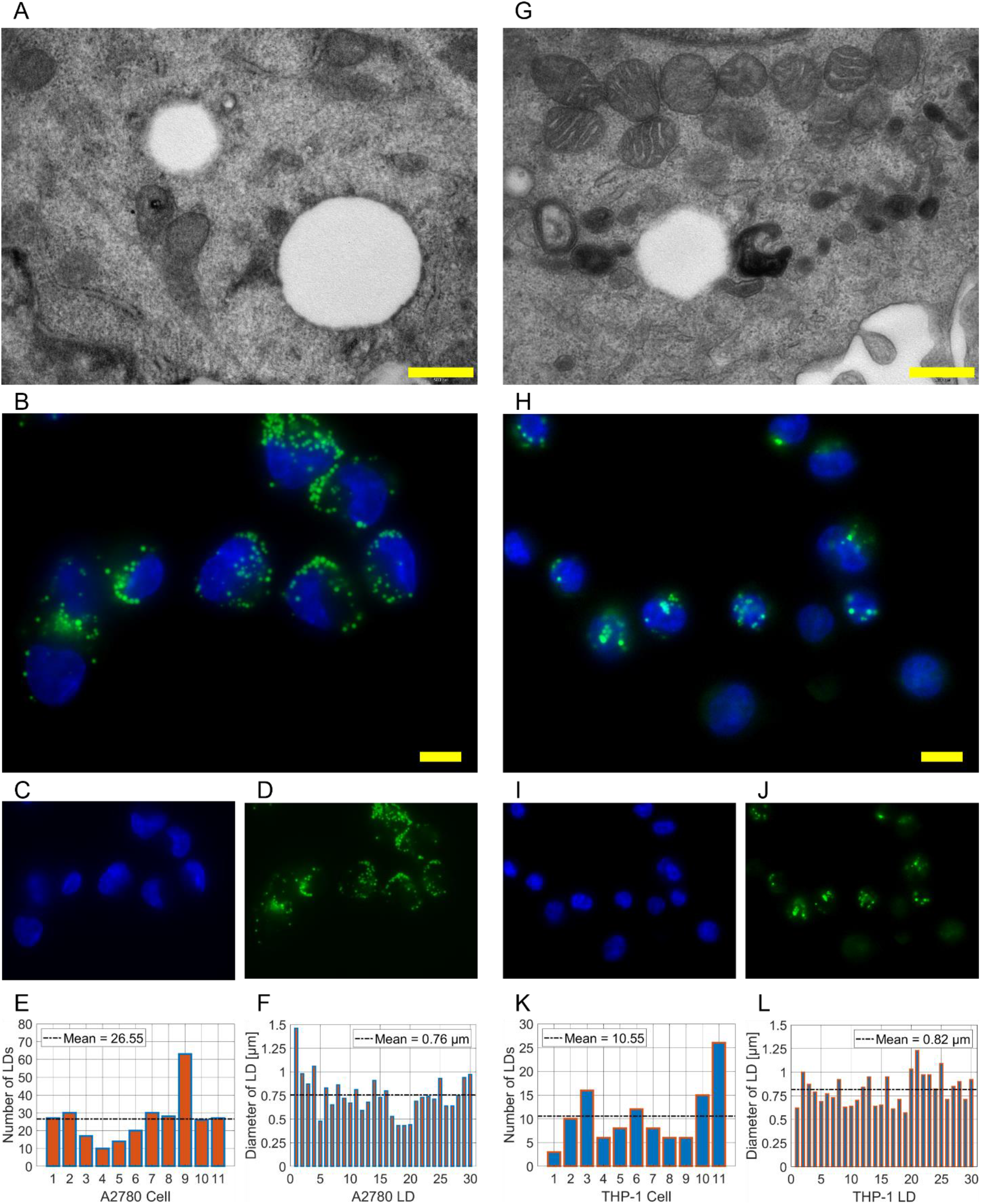
Conventional 2D imaging of A2780 (A-F) and THP-1 (G-L) live cells. (A,G) TEM images of LDs. Scale bar is 0.5 μm. (B,H) Representative FM images of nuclei (blue) and LDs (green). Scale bar is 10 μm. (C,I) FM images of nuclei stained with Hoechst. (D,J) FM images of LDs stained with Nile Red. (E,K) Number of LDs in 11 live cells imaged by FM. (F,L) Diameters of 30 LDs imaged by FM.

## Discussion

In the last few years, LDs have been demonstrated to be involved in increasing number of physiological processes, due to both their own activity and the way they relate to the other intracellular organelles. In particular, LDs dysregulation has been found in several pathologies. For this reason, imaging systems able to provide a reliable analysis about LDs are strongly requested to deepen these aspects and to promote the spread of diagnostic or therapeutical tools respectively based, e.g. on the detection of LDs as biomarkers or the regulation of their activity. The high spatial resolution and contrast of TEM make it a gold-standard method for highly detailed imaging of these organelles. However, the advantages of electronic microscopy are counterbalanced by the limitation of imaging 2D ultrathin slices (^~^ 80 nm) with a very small FOV (a few square microns). In fact, in a cell with a diameter of tens of microns, TEM imaging is not suitable for quantitative counts, since the measurements are often underestimated and, in turn, LDs can be found only in a small percentage of the total number of analysed cells. Moreover, sample preparation requires difficult and time-consuming protocols thus preventing statistical studies on a large number of cells. Along with TEM, FM is commonly employed to study LDs inside cells, even though this kind of 2D imaging is limited by phototoxicity, which can damage the sample, and by photobleaching, which hinders long time experiments. Moreover, only a few properties can be measured from the FM images, like the number or the size of LDs, with quantification of these parameters in a high number of cells being time consuming. It is important to remark that both TEM and FM are 2D imaging techniques, which provide only limited information about LDs in the absence of their 3D representation. A possible solution is 3D confocal microscopy, in which however fluorescent labelling is required, while the TPM is a label-free technique. Recently TPM has been demonstrated to be a powerful tool for the stain-free quantification of 3D features related to the cell biophysical properties. In particular, TPM has been already applied for LDs analysis (28–33). However, so far TPM has been realized only in static environment for studying adhered cells, thus not overcoming the low-throughput issue also typical of 3D confocal microscopy. Instead, here for the first time we have proposed the application of the TPM technique in flow-cytometry conditions for the 3D visualization and full-characterization of LDs inside stain-free suspended cells. In particular, we have demonstrated the reliability of the in-flow TPM system by reconstructing the 3D RI distribution of LDs inside 54 A2780 human ovarian cancer cells and 34 THP-1 monocyte cells. It is important to note that in our approach we decided to set the resolution of the DH system limited at 0.5 μm in order to privilege instead a larger FOV (~ 0.41 mm2), allowing imaging tens of cells in each hologram of the recorded sequence. For this reason, we limit our capability to detect and visualize LDs having size lower than 1 μm. Nevertheless, the DH TPM set up can be easily adjusted with a different microscope objective offering higher resolution. Thus, the resolution limit in our tomograms is not an intrinsic limitation of DH TPM but rather a constraint imposed by our specific optical design for the reported experiments. On the other hand, the DH capability of providing QPMs with nanometric resolution has already been demonstrated (49). Beyond features like the volume or the RI-based statistics that can be accessed also by the static TPM, we have also quantified the 3D spatial localization of LDs inside suspended cells, which is fundamental to study the still unknown relations between LDs and other intracellular organelles in physiological and disease contexts. Interestingly, despite LDs are usually expected to be found adjacent to the ER (45), our analysis revealed LDs assembled mainly near the nucleus in A2780 model, which may possibly reflect their particular function in ovarian cancer cells. Another important advantage of our in-flow TPM approach is the high-throughput property, since it can be potentially exploited to perform a statistical analysis on thousands of cells per experiment. Above all, we want to stress that the flow-cytometry modality is ideal for reproducing the physiological conditions of certain cell types, e.g. white blood cells or circulating tumour cells, thus boosting the TPM potential for diagnostic applications like liquid biopsy (50).

## Materials and Methods

### Sample preparation

For FM experiments, human ovarian cancer cell line A2780 was purchased from Sigma Aldrich (#93112519), whereas THP-1 monocyte cell line was supplied by a third part. Both cell lines were cultured in RPMI 1640 Medium (Euroclone #ECB9006L), supplemented with 10% FBS (Euroclone #ECS5000L), 2mM L-Glutamine (Euroclone #ECB3000D) and 1% Penicillin/Streptomycin (Euroclone #ECB3001D), and maintained at 37°C in a humidified atmosphere with 5% CO2. A2780 were grown in a standard adherent culture on a 2D surface, whereas THP-1 cell line was grown in suspension. EVOS M5000 Imaging System (ThermoFisher Scientific #AMF5000) was used for cell line monitoring.

For in-flow TPM experiments, the THP-1 monocyte cell line was grown in suspension and cultured in RPMI 1640 Medium (Life technologies, ref 31870-025), supplemented with 10% FBS (Life Technologies 10270), 2mM L-Glutamine (Lonza, Cat N.: BE17-605E) and 1% Penicillin/Streptomycin (Lonza, Cat N. DE17-602E), and maintained in cell culture flask (Corning, product number 353018) at 37°C in a humidified atmosphere with 5% CO2. The day of the experiment they were harvested from the cell culture flask and centrifuged for 5 min at 1500 rpm, resuspended in complete medium and injected into the microfluidic channel at final concentration of 3×105 cells/mL. The A2780 cell line was grown as a monolayer and cultured in RPMI 1640 Medium (Life technologies 31870-025) supplemented with 10% FBS (Life Technologies 10270), 1% Penicillin/Streptomycin (Life Technologies 15070-063) and 2mM L-Glutamine (Lonza BE 17-605E). They were cultivated at 37°C in an incubator with 5% CO2. Later, cells were washed twice with PBS 1x (Life technologies 10010023) and incubated with 1.5mL of 0.05% trypsin–EDTA solution (Sigma, T4049-100ML) for 5 min. The cells were resuspended with a solution composed by PBS plus 10% FBS, to neutralize the trypsin effect, and viability was assessed through Trypan Blue solution 0.4% (Sigma T8154) according to manufacturer’s procedures. Then, the cells were injected into the microfluidic channel at final concentration of 5×105 cells/mL.

### TEM imaging

THP-1 and A2780 cells were seeded at the density of 2×105 cells/well and cultured in a 6-well plate. After washing, the cells were fixed in 2.5% buffered glutaraldehyde in the culture plate for 20 min at room temperature, scraped, centrifuged for pellet recovery, and stored overnight at 4°C. After post-fixation in 1% buffered osmium tetroxide for 2 h at 4°C, samples were rinsed, dehydrated in increasing concentrations of ethanol, embedded in epoxy resin and sectioned by means of ultramicrotome. Thick sections (80 nm) were stained with uranyl acetate followed by lead citrate and examined in a Philips CM100 (FEI Company, ThermoFisher, Waltham, MA, USA) Transmission Electron Microscope. Images were taken with an Olympus Megaview II digital camera integrated with iTEM image processing software.

### FM imaging

A2780 (1.5×105 cells/mL) were seeded on coverslips and cultured for 24 h. The cells were then washed twice with PBS and incubated with 5 μM Nile Red (Sigma, N3013) and 1 μg/mL Hoechst 33342 (Invitrogen, H1399) in RPMI 1640 complete medium for 30 min. THP-1 (1.5×105 cells/mL) were seeded in suspension and after 24 h they were pelleted, washed twice with PBS and incubated for 30 min in RPMI 1640 complete medium containing Nile Red and Hoechst at above indicated concentrations. For both cell lines, after staining, the medium was discarded and substituted with complete medium without phenol red. For imaging, THP-1 cells were placed on a microscope slide and “coverslipped”. LDs and nuclei were acquired in live cells with a digital imaging system, using an inverted epifluorescence microscope with ×63/1.4 numerical aperture (NA) oil objective (Nikon Eclipse Ti-U, Nikon), applying 500 ms and 100 ms exposure time, for Nile Red and Hoechst, respectively. Images were captured with a back-illuminated Photometrics Cascade CCD camera system (Roper Scientific) and elaborated with Metamorph Acquisition/Analysis Software (Universal Imaging Corp). The number and the dimension of LDs were quantified using ImageJ software (51). Statistical analysis comparing A2780 and THP-1 cells was performed by applying a two-tailed Student’s T-test.

### In-flow TPM imaging

The DH microscope is developed in off-axis telecentric configuration employing a Mach-Zehnder interferometric scheme, as sketched in Fig. 6A. A laser light (Laser Quantum Torus 532) emitting at 532 nm with an output power equal to 750 mW is used as illumination source. The laser beam is split by a polarizing beam splitter (PBS) cube which separates it in object and reference beams, the latter is transmitted and the former is reflected. In addition, to balance the ratio between intensity of object and reference beam, maintaining the same polarization, two half-wave plates (HWP) are placed in front of and behind the PBS, respectively. Object beam passes through a microfluidic chip where cells flow and rotate and then it is collected by a microscope objective (MO1 Zeiss Plan-Apochromat 40× NA=1.3 Oil immersion) and sent to a tube lens (TL1 with focal length 150 mm). Reference beam passes through a beam expander shaped by a microscope Objective (MO2 NewPort 20× NA=0.40) and a second tube lens (TL2 with focal length is 250 mm). Then, both collimated beams are recombined by a beam splitter cube (BS) with a small angle between them in order to achieve off-axis configuration and generate interference pattern digitally recorded by the CMOS camera (Genie Nano-CXP Cameras 5120×5120 pixels, until to 80 fps and pixel size equal to 4.5 μm). The FOV is 640×640 μm2 with a spatial resolution of 0.5 μm. The camera is equipped by a video recording system that ensure long time acquisition mode.

**Fig. 6.**
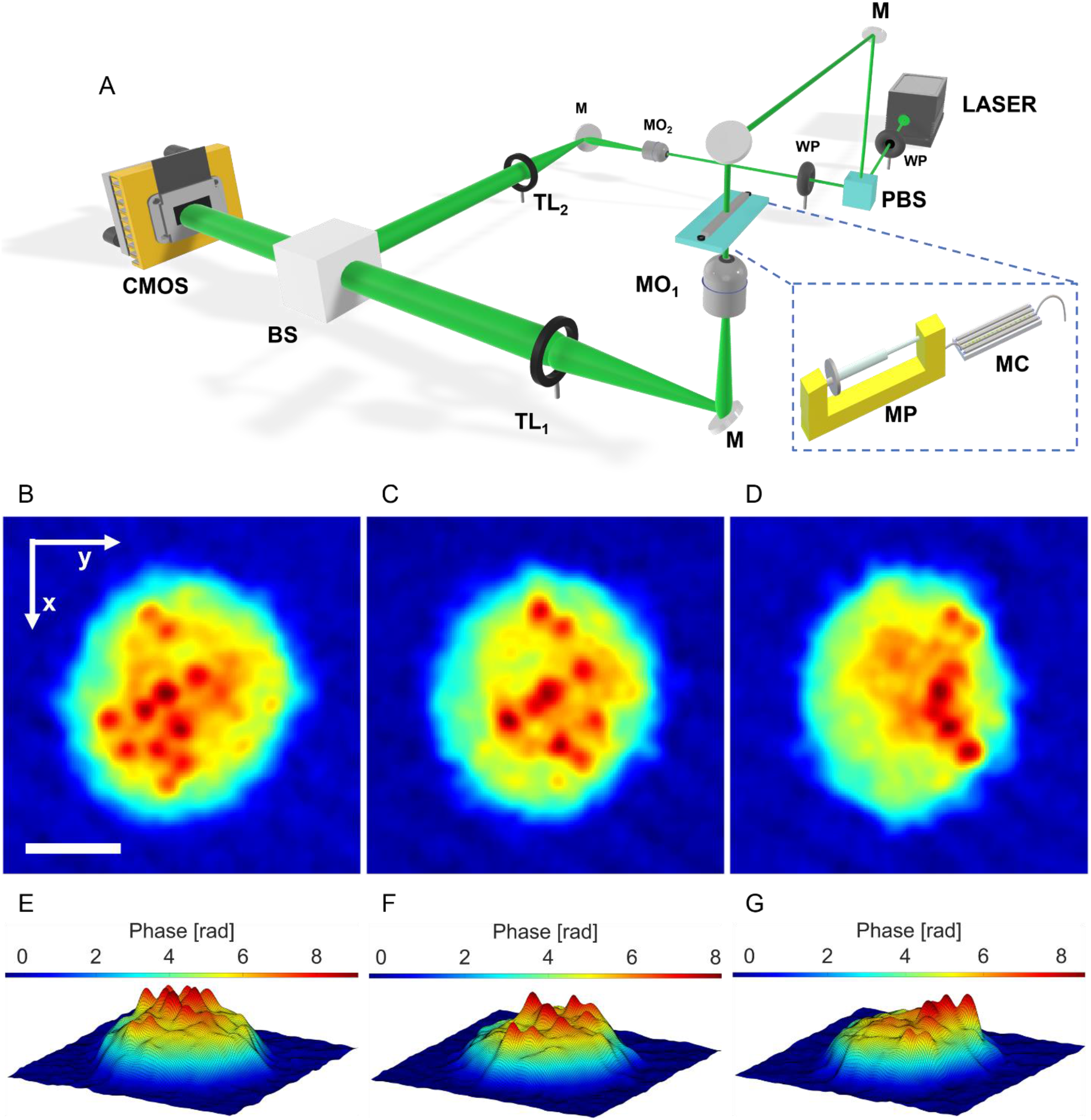
In-flow TPM system. (A) DH microscope in off-axis configuration. PBS – Polarizing Beam Splitter; WP –Wave Plate; M – Mirror; L1, L2 – Lens; MO – Microscope Objective; MC – Microfluidic Channel; MP – Microfluidic Pump; TL – Tube Lens; BS – Beam Splitter; CMOS – Camera. (B-D) Three QPMs of an A2780 cell while flowing along the y-axis and rotating around the x-axis. The spots with the biggest phase values (dark red) are the LDs. Scale bar is 5 μm. (E-G) Pseudo-3D visualization of the 2D QPMs in (B-D), respectively, in which the LDs are well-separated from the outer cell because of their greater height.

Sample is prepared and inserted in the automatic syringe pump (Syringe Pump neMESYS 290N), that is a low-pressure system which allows a high-precision and pulsation-free dosing of liquids at micro and nanoliter scale ensuring a very homogeneous flow inside the microchannel (Microfluidic ChipShop 10000107). In a typical operation mode, holographic video sequences are registered at 30 fps at a flow rate of about 50 nl/s that allow the 360° rotation of the flowing cells in a FOV. In fact, due to the parabolic velocity profile related to the laminar flow generated by the microfluidic pump, the cells rotate around the x-axis while flowing in the microfluidic channel along the y-axis, thus allowing the recording along the z-axis of digital holograms of the same single cell at multiple viewing angles (34, 35). For each hologram of the sequence, the corresponding QPM is numerically retrieved. In particular, the hologram is demodulated to select the real diffraction order (52). A region of interest (ROI) is selected around the analysed cell, and then it is propagated at different distances along the optical axis in order to refocus it by minimizing an image contrast metric (Tamura Coefficient) computed on the amplitude part of the complex wavefront (53). The phase aberration is compensated by subtracting a reference hologram acquired without the sample in the imaged FOV (54). The phase component of the refocused complex wavefront is finally denoised (55) and unwrapped (56) to obtain the QPM. In order to avoid motion artefacts in the tomographic reconstructions, each ROI is centred with respect to the cell centroid. Three typical QPMs of an A2780 cell are reported in Figs. 6B-D, in which the rotation around the x-axis of the cell flowing along the y-axis can be observed. Moreover, as the RIs of the LDs are higher than the other organelles, they can be recognized within the QPMs as the red spots. To mark this property, we show the same QPMs in a pseudo-3D visualization in Figs. 6E-G, in which the height codifies the phase values. Hence, LDs correspond to the isolated peaks, which change their position because of the cell rotation in the microfluidic channel. The in-flow TPC allows overcoming most of the main drawbacks of the conventional TPM techniques (34), which are based on the scanning along different beam direction if the sample is fixed or along a unique beam direction if the sample is rotated by mechanical/optical forces.

In fact, the full rotation of the flowing samples induced by controlled hydrodynamic forces allows recording in high-throughput modality (hundreds of flowing and rotating cells per minutes) a complete phase information all around them, without introducing external cell perturbations. However, to ensure these advantages, the rolling angles are not known a priori, although they are requested to perform the tomographic reconstruction. Recently, we proposed a rolling angles recovery method to solve this issue, based on the identification of phase similarities (36). The presence of intracellular LDs provides a great help in making more accurate the rolling angles recovery and then the 3D RI tomogram (see the Supplementary Materials). Once the unknown viewing/rolling angles are estimated, the pairs consisting of the QPMs and the corresponding rolling angles are given in input to the Filtered Back Projection algorithm (57), in order to reconstruct the 3D RI spatial distribution at the single cell level. A typical in-flow TPM experiment takes few minutes (2-3 min) for recording the holographic sequences containing hundreds of flowing/rolling cells. In particular, on average the holographic sequence of each flowing/rolling cell is made of 200-300 images, each corresponding to a different viewing angle. The ex-post numerical process to reconstruct the 3D RI tomogram requested about 10-20 min, even though this time can be drastically reduced up to few minutes through a deep-learning approach.

## Acknowledgments

The THP-1 monocyte cell line was a kind gift from Dr. Ilaria Malanchi from The Francis Crick Institute, London, UK.

## Author contributions

I.K., P.M., L.M. and P.F. conceptualized the research; D.S. and L.M. set the holographic tomography system and were responsible of the holographic acquisitions; D.S., V.B., D.P., L.M. and P.M. contributed to optimize the holographic acquisitions and data preparation; M.M. and D.d.G. prepared the biological samples and contributed to the holographic experiments; D.P. and P.M. were in charge for the holographic reconstructions and tomograms; D.P., P.M., V.B., L.M. and P.F. analyzed and discussed the tomographic reconstructions and data. I.K., S.L. and L.I. performed FM experiments and data analysis; G.P. and S.V. performed the TEM; all the authors contributed to critical discussion of the results and contributed to write the manuscript. P.F. supervised the research.

## Data and materials availability

All data are available in the main text or the supplementary materials.

## Supplementary Materials for

### Rolling angle recovery

To recover the unknown rolling angles of the flowing cell, a full rotation is detected through a suitable phase image similarity metric, and then a proportion with the positions along the flow direction is implemented (36). However, this method suffers when the similarity metric does not have a trend with pronounced minima. In that case, an error can be committed in retrieving the unknown rolling angles, which propagates to the tomographic reconstruction. Instead, the presence of intracellular LDs provides a great help in making more accurate the rolling angles recovery and then the 3D RI tomogram. In fact, as shown in Figure 6(B-G), the LDs are phase peaks which move in the QPM sequence according to the cell rotation, thus providing a highly distinguishable marker for the phase similarity. This property is highlighted in the phase similarity metric reported as example in Fig. S1A, regarding the QPM sequence of the A2780 cell analyzed in Fig. 6(B-G). It is computed by comparing all the QPMs with the first one of the sequence. Hence, it is 0 in the first frame, while the other local minima, which are well defined thanks to the LDs, correspond to full rotations (i.e., 360°). Indeed, a full rotation can be easily detected when LDs come back to their starting positions, as shown in Figs. 4B,C.

**Fig. S1.**
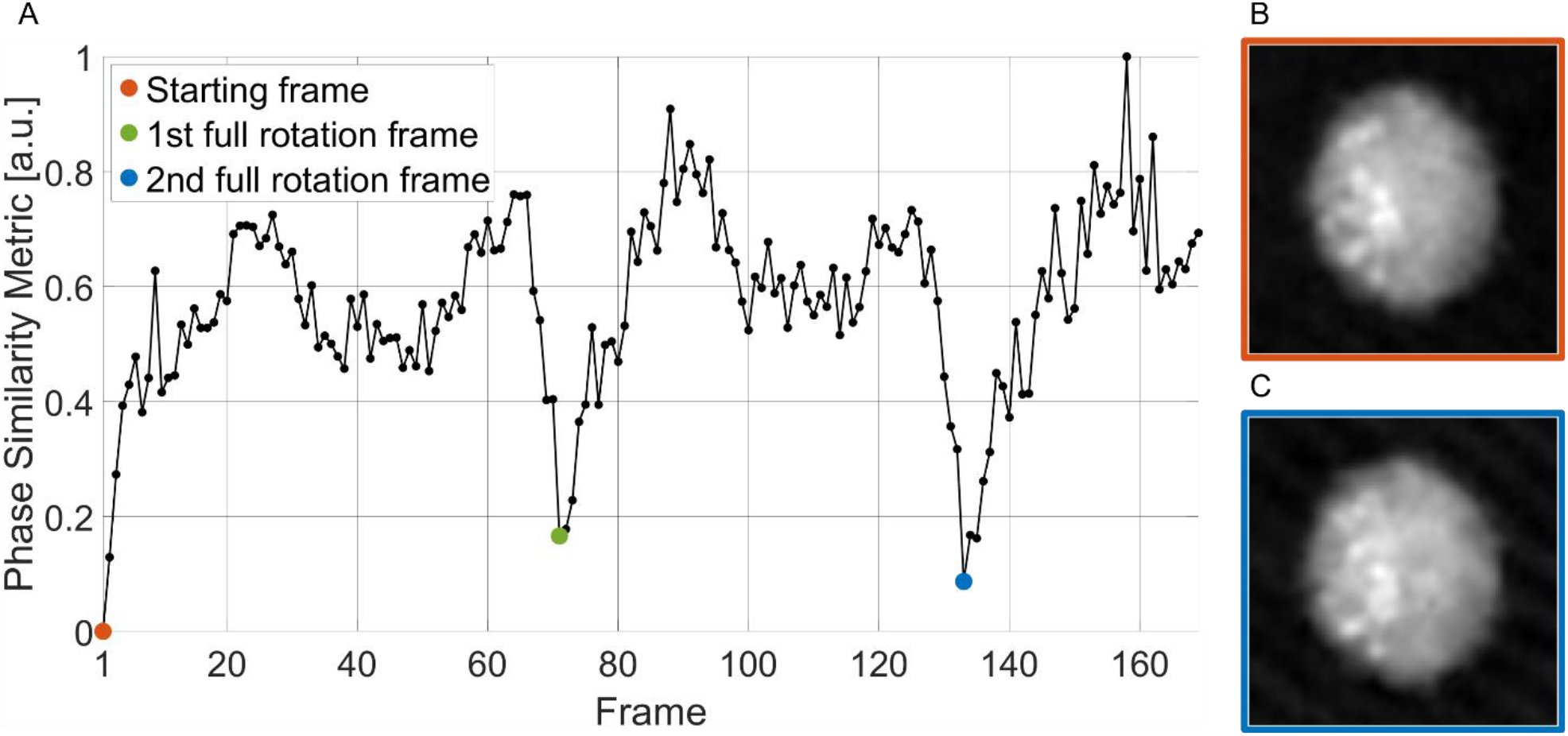
LDs-aided method for the rolling angles recovery in an A2780 live cell. (**A**) Trend of the phase similarity metric, which is null in the starting frame of the QPM sequence (orange dot) and is minimum when the first (green dot) and the second (blue dot) full cell rotations have occurred. (**B**) QPM at the first frame and (**C**) QPM after two full rotations, in which LDs are located in the same positions.

**Table S1.** Average values and standard deviations of the features about each LD segmented in 54 A2780 and 34 THP-1 live cells.

|  |  |  |
| --- | --- | --- |
|  | *A2780* | *THP-1* |
| *RI Mean Value* | $1.437 \pm 0.006$ | $1.420 \pm 0.004$ |
| *RI Standard Deviation* | $0.008 \pm 0.003$ | $0.006 \pm 0.002$ |
| *RI Entropy* | $2.851 \pm 0.570$ | $2.389 \pm 0.501$ |
| *RI Kurtosis* | $2.310 \pm 0.280$ | $2.236 \pm 0.238$ |
| *RI Skewness* | $0.520 \pm 0.134$ | $0.479 \pm 0.131$ |
| *Volume [$\mu\text{m}^3$]* | $2.267 \pm 2.430$ | $1.678 \pm 1.272$ |
| *Equivalent Radius [$\mu\text{m}$]* | $0.751 \pm 0.215$ | $0.708 \pm 0.141$ |
| *Surface Area [$\mu\text{m}^2$]* | $8.873 \pm 7.038$ | $7.109 \pm 3.672$ |
| *Sphericity [a.u.]* | $0.914 \pm 0.086$ | $0.940 \pm 0.074$ |
| *Dry Mass [pg]* | $1.757 \pm 1.920$ | $1.085 \pm 0.861$ |
| *LD-Cell Normalized Distance [a.u.]* | $0.666 \pm 0.178$ | $0.686 \pm 0.139$ |
| *LD-LDs Centroid Normalized Distance [a.u.]* | $0.251 \pm 0.093$ | $0.261 \pm 0.104$ |
| *1<sup>st</sup> Order Coefficient Parabolic Fitting [$\mu\text{m}^{-1}$]* | $-0.037 \pm 0.011$ | $-0.028 \pm 0.007$ |
| *2<sup>nd</sup> Order Coefficient Parabolic Fitting [$\mu\text{m}^{-2}$]* | $-0.006 \pm 0.010$ | $-0.003 \pm 0.004$ |

